# A dataset of adult heart and liver mass after placental *Insulin-like growth factor 1* overexpression and insufficiency in mice

**DOI:** 10.1101/2025.05.23.655797

**Authors:** Faith M Fairbairn, Annemarie J Carver, Robert J Taylor, Hanna E Stevens

## Abstract

The placenta is an important producer of hormones essential for fetal development. Insulin-like growth factor 1 (IGF1) is a hormone primarily produced in the placenta in utero and is an important regulator of various developmental pathways including those in heart and liver. Embryonic disruptions in these developmental pathways can lead to lifelong changes and are often associated with chronic disease. Further, the placenta has sex-specific impacts on offspring development in response to hormonal changes. Previous work has shown that altered expression of *Igf1* in the placenta results in sexually dimorphic changes to placental and fetal developmental outcomes. Here, mice underwent placental-targeted CRISPR manipulation for overexpression or insufficiency of *Igf1*. At the time of euthanasia, heart and liver tissues were collected and weighed. This dataset presents the heart and liver mass of these postnatal mice. There was a significant increase in proportional heart mass in placental *Igf1* overexpression adult female mice and a trending increase in proportional liver mass in placental *Igf1* overexpression adult male mice. No significant changes in heart or liver mass were seen in placental *Igf1* insufficiency mice. These data provide insight into the impact of placental IGF1 on long-term heart and liver development.

**VALUE OF THE DATA:**

- There is significant evidence for the role of early genetic changes in influencing long-term health outcomes, as laid out by the Developmental Origins of Health and Disease (DOHaD) hypothesis [1]. According to this hypothesis, genetic factors may be critical in determining the timing and severity of chronic disease, with varying effects based on sex. Genetics of the placenta, which makes up the maternal-fetal interface, plays an important role in modulating exposures associated with the DOHaD hypothesis [2].
- The placenta provides essential hormones to the fetus during pregnancy [3]. Placental changes are associated with the development of chronic disease and metabolic changes [4,5]. Disruptions in placental functions have been linked to defects including congenital heart disease which affects approximately 40,000 babies each year in the United States [6,7]. The placenta is also linked to metabolic diseases later in life such as nonalcoholic fatty liver disease, a chronic liver disease which has increased in prevalence by over 50% from 1990 to 2019 [5,8,9].
- Insulin-like growth factor 1 (IGF1) is a placentally produced factor that regulates pathways involved in fetal growth and development and has been shown to be critical in growth of the heart and liver [10-13]. Despite the importance of the placenta and IGF1 in heart and liver growth, specific links between placental *Igf1* expression and developmental outcomes remain understudied.
- Placental function is known to have sex-specific impacts on fetal growth [14]. Further, *Igf1* expression in the placenta is linked to differences in offspring developmental outcomes by sex [15]. Placental *Igf1* overexpression and insufficiency affect offspring in a sexually dimorphic manner. IGF1 is a hormone and interacts with sex hormones, likely contributing to sex differences in response to changes in *Igf1* expression [16]. Further research, including the work done to produce this dataset, may help clarify the role of placenta *Igf1* expression in fetal outcomes, specifically regarding sex differences.
- The data presented in this paper provide insight into the effects of placental Insulin-like growth factor 1 overexpression and insufficiency on adult heart and liver mass. More research is needed to understand specific functional impacts on these organs. Further, understanding the effects of placental genetic changes may support the development of future treatments and therapies for placental insufficiencies.

## BACKGROUND

The placenta produces and regulates essential hormones involved in growth signaling [3]. Previous studies have shown that placental manipulations influence offspring development [15,17]. The placenta is the primary source of Insulin-like growth factor 1 (IGF1) during fetal development in humans and mice [10,11]. Sex-specific placental function and *Igf1* expression have a role in sexual dimorphisms of offspring outcomes [14,15]. IGF1 supports placental morphogenesis and function and regulating overall fetal development, with specific impacts on heart and liver [10,12,13].

Placenta and heart share important developmental pathways, with placental abnormalities linked to congenital heart disease [6]. IGF1 supports cardiomyocyte growth in gestation and is linked to cardiac hypertrophy [12]. Placental function is also linked to hepatocyte proliferation [13]. Placental morphological and hormonal variations, including IGF1, are implicated in metabolic diseases often involving the liver [5].

Here, placental *Igf1* overexpression (Igf1-OE) or *Igf1* insufficiency (Igf1-KO) was induced in mice on embryonic day 12 (E12) via CRISPR placental manipulation [17]. Previously, Igf1-OE and Igf1-KO were differentially induced in males and females and resulting impacts on placenta and offspring brain were sex-specific. [15]. Therefore, placental *Igf1* changes could influence other offspring outcomes sex-specifically, namely heart and liver growth, for which data are presented here.

## DATA DESCRIPTION

The dataset includes adult mouse Igf1-OE and Igf1-KO offspring values for mass of heart and liver compared to appropriate controls, ConAct and ConCas9, respectively. Here, the data are presented as raw values for mass and also adjusted as a proportion of body mass. There were no significant changes in adult body mass in either Igf1-OE or Igf1-KO adult mice compared to controls (data not shown) [15].

Effects of placental *Igf1* manipulation on heart mass for adult Igf1-OE and Igf1-KO mice are shown in Figure 1. Adult female Igf1-OE mice had significantly increased heart mass (13.06%) and proportional heart mass (10.31%) compared to the appropriate control group (Figure 1A,B), with no change for adult male Igf1-OE mice (Figure 1C,D). Placental Igf1-KO had no significant impact on heart mass in adult mice (Figure 1E-H).

**Fig 1.**
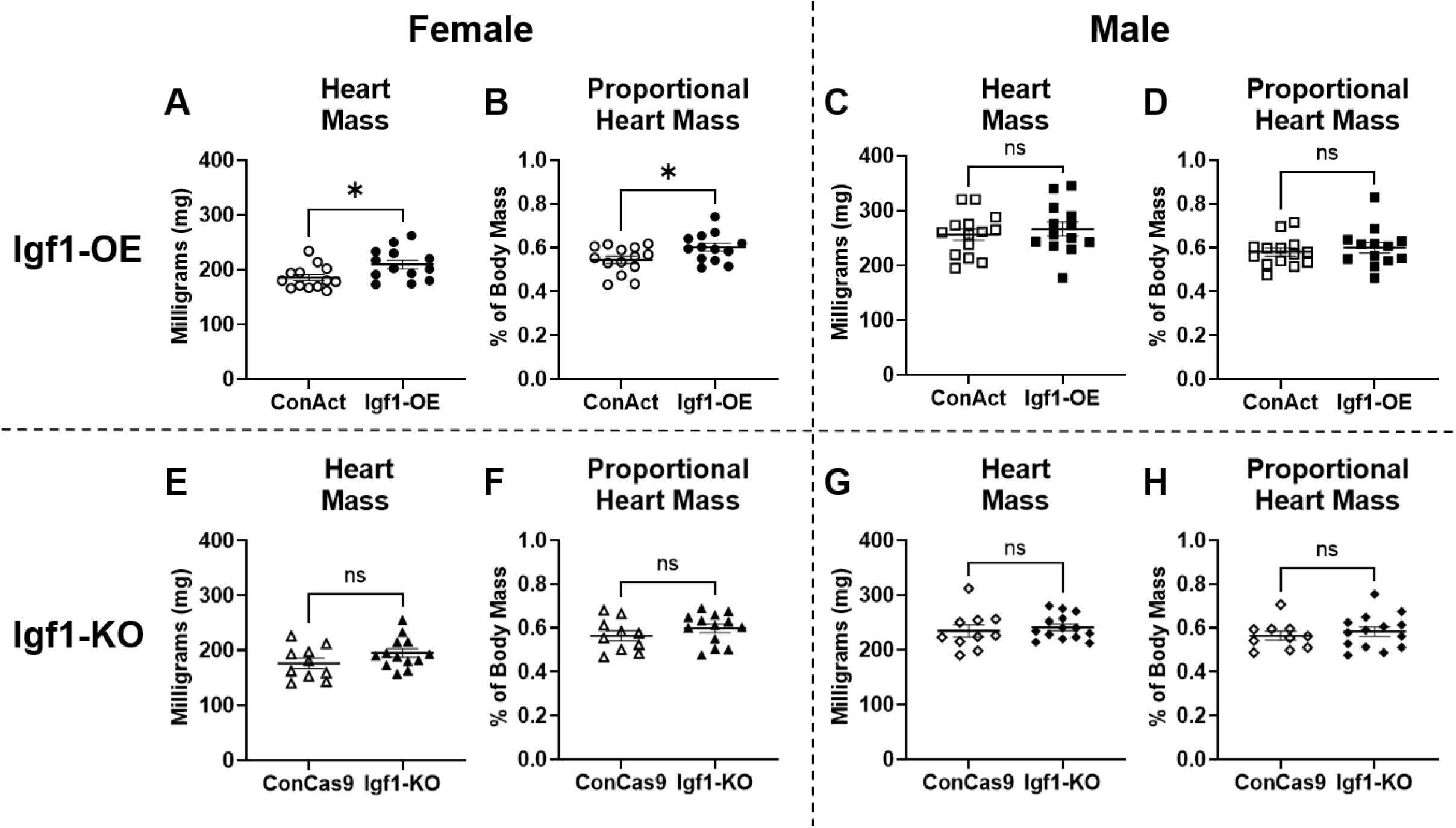
Heart mass and heart mass as a proportion of body mass separated by sex in adult Igf1-OE mice compared to control (ConAct) (A-D). Significant increase in Igf1-OE female heart mass and proportional heart mass (p=0.0226 and 0.0324 respectively). Heart mass and heart mass as a proportion of body mass separated by sex in adult Igf1-KO mice compared to control (ConCas9) (E-H).

In Figure 2, liver mass for adult Igf1-OE and Igf1-KO mice are displayed. While females showed no significant increase, liver mass averages across groups showed similar percent increases as for the heart (15.27% for mass and 8.71% for proportional mass) (Figure 2A,B). Interestingly, adult male Igf1-OE mice had a trending increase in proportional liver mass (4.53%) (Figure 2C,D). Placental Igf1-KO had no significant impact on liver mass in adult mice (Figure 2E-H).

**Fig 2.**
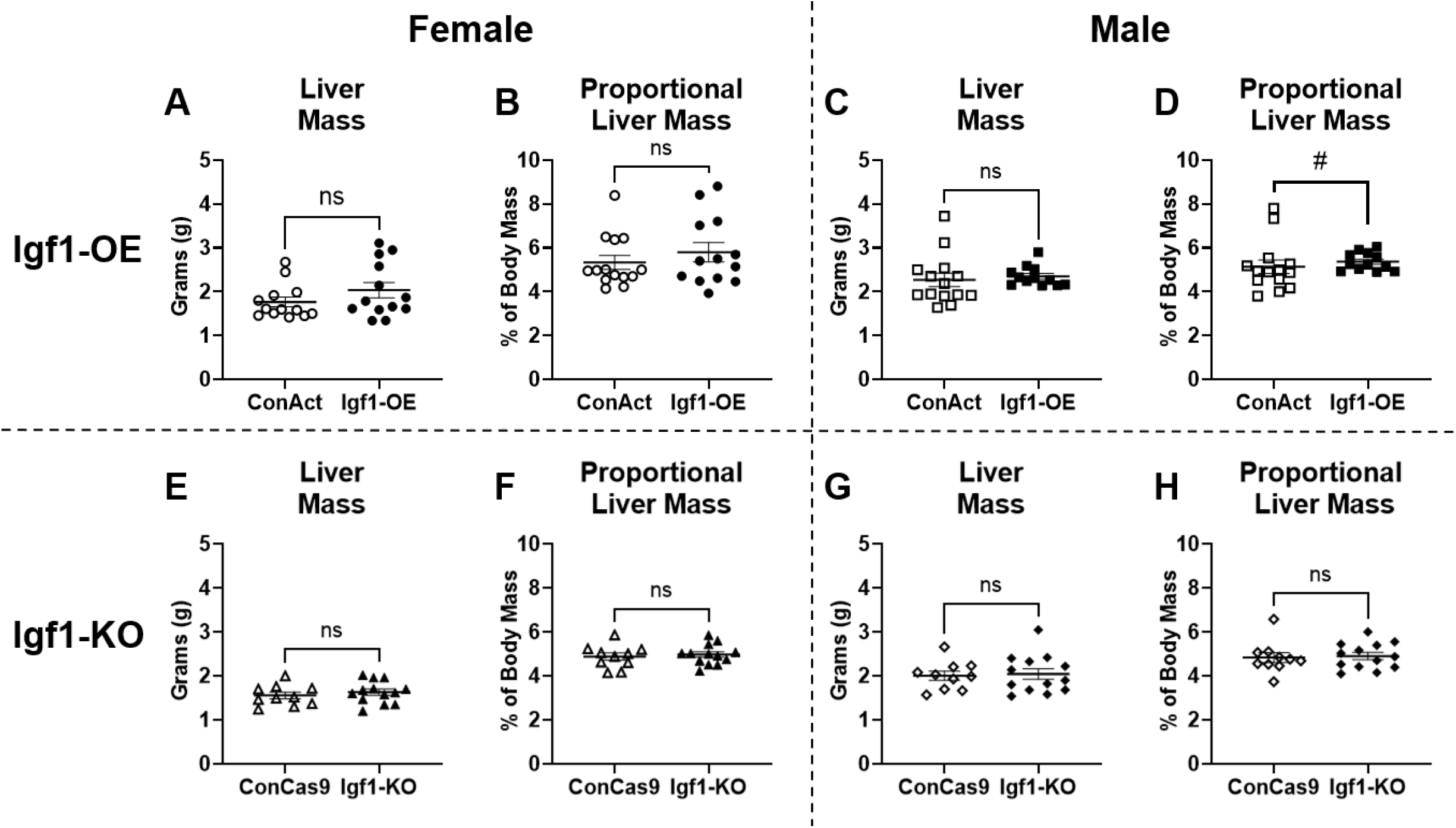
Liver mass and liver mass as a proportion of body mass by sex in adult Igf1-OE mice compared to control (ConAct) (A-D). Trending increase in proportional liver mass in adult Igf1-OE male mice (p=0.0757). Liver mass and liver mass as a proportion of body mass by sex in adult Igf1-KO mice compared to control (ConCas9) (E-H).

## EXPERIMENTAL DESIGN, MATERIALS AND METHODS

### Animal handling and placental-targeted manipulation

All experiments were done with Institutional Animal Care and Use Committee approval from University of Iowa. All mice used in this study were on 12-hour light cycle and given food and water ad libitum. Female CD-1 mice (Charles River, strain 022, 2-4 months old) were bred with CD-1 males until observation of copulation plug, designated as embryonic day 0 (E0). Females were then singly housed until E12, where they underwent placental-targeted CRISPR manipulation [17]. E12 dams were anesthetized and surgical plane maintained with isoflurane, abdominal incisions made, uterine horns exposed, and each placenta within the litter was independently injected with either IGF-I CRISPR Activation Plasmid (m) (sc-421056-ACT) referred to as (Igf1-OE), IGF-I CRISPR/Cas9 KO Plasmid (m) (sc-421056) referred to as (Igf1-KO), Control CRISPR Activation Plasmid (sc-437275) (ConAct), or Control CRISPR/Cas9 plasmid (sc-418922) (ConCas9) (All plasmids from Santa Cruz Biotechnology). Here, Igf1-KO is hypomorphic with *Igf1* expression knocked out in some but not all placental cells resulting in an insufficiency. Following each injection, each placenta was electroporated. Then, uterine horns were returned, incisions sutured, and the dam allowed to recover and give birth. Offspring were weaned on postnatal day 21 into sex-specific group-housing.

### Tissue collection

12-to 14-week-old offspring were deeply anesthetized with ketamine/xylazine cocktail and transcardially perfused with PBS followed by 4% paraformaldehyde. Body mass was recorded prior to perfusion. Liver and heart were then removed with consistent dissection methods and immediately weighed.

### Data Analysis

Heart and liver mass as well as percentage of heart and liver mass to total body mass were compared separately by sex and to the appropriate control group for CRISPR plasmid used. Outliers were determined in GraphPad prism by the ROUT outlier test. After exclusion of outliers, all comparisons were performed on GraphPad Prism using an unpaired Welch’s t-test. Significance was defined as *p < 0.05 and a trend was defined as #p<0.1.

## LIMITATIONS

This project used CRISPR manipulation to model increased and decreased *Igf1* expression in the placenta to understand impacts on heart and liver growth. It is important to note that this manipulation creates an artificial change in the *in utero* environment and, while a model of expression changes, may not be representative of a physiological range of *Igf1* expression levels. Additionally, the CRISPR manipulation occurred on embryonic day 12, and is, therefore, not indicative of placental *Igf1* expression changes throughout an entire pregnancy or beginning at other time points during pregnancy. The weights displayed in this dataset were collected at adult time points and are, therefore, not reflective of any potential earlier changes that may have normalized by the time the tissue was collected. There was no indication that the induced changes had negative impacts on the overall success of the pregnancy or offspring viability.

## ETHICS STATEMENT

This work involved animal experiments and complied with the National Institutes of Health guide for the care and use of laboratory animals (NIH Publications No. 8023, revised 1978). Mouse care, handling, and husbandry were performed as approved by the University of Iowa IACUC.

## CRediT AUTHOR STATEMENT

**Faith Fairbairn:** Investigation, Visualization, Writing—original draft, Writing—reviewing and editing. **Anna Carver:** Conceptualization, Data curation, Formal analysis, Funding acquisition, Investigation, Methodology, Project administration, Supervision, Validation, Visualization, Writing—original draft, Writing—reviewing and editing. **Robert Taylor:** Conceptualization, Investigation, Methodology, Validation, Writing—original draft, Writing—reviewing and editing. **Hanna Stevens:** Conceptualization, Funding Acquisition, Methodology, Project administration, Resources, Supervision, Writing—reviewing and editing.

## ACKNOWLEDGEMENTS

The authors would like to acknowledge the following funding sources: R01 MH122435 (H.E.S), NIH T32GM008629 (A.J.C.), and NIH T32GM145441 (A.J.C.) as well as the University of Iowa Graduate College (A.J.C).

